# Predator phylogenetic diversity decreases predation rate via antagonistic interactions

**DOI:** 10.1101/089144

**Authors:** A. Andrew M. MacDonald, Gustavo Q. Romero, Diane S. Srivastava

## Abstract

**Background:** Predator assemblages can differ substantially in their top-down effects on community composition and ecosystem function, but few studies have sought to explain this variation in terms of the phylogenetic distance between predators. The effect of a local predator assemblage will depend on three things – which predators tend to co-occur, how similar their prey preferences are, and how they interact with each other and the whole community. Phylogenetic distance between predators may correlate with each of these processes, either because related predators are more likely to share the same traits, and therefore have similar habitat and prey preferences, or because predators are more likely to compete, and therefore diverge in habitat and prey preferences. Therefore, the phylogenetic structure of predator assemblages could provide a unifying framework for predicting how predators will impact their prey - and therefore any ecosystem functions mediated by their prey.

**Methods:** We examined the effects of predators on macroinvertebrate food webs found in bromeliads, combining field observations, laboratory feeding trials and a manipulative experiment. We determined whether the phylogenetic distance between predators could explain: the co-observed occurrence of predator species among bromeliads, overlap in prey preferences under no-choice conditions, and effects of predator composition on prey survival, prey composition and ecosystem processes.

**Results:** We found that phylogenetic distance does not predict either the co-occurrence of predator species nor the overlap in their prey preferences. However, our manipulative experiment showed that prey mortality decreased as the phylogenetic distance between predator species increased, reflecting antagonistic interactions among more distant predators. These effects of phylogenetic distance on prey mortality did not translate into effects on ecosystem function, as measured by rates of detrital decomposition and nitrogen cycling.

**Discussion:** We conclude that the effects of predator phylogenetic diversity on the bromeliad food web are primarily determined by antagonistic predator-predator interactions, rather than habitat distribution or diet overlap. This study illustrates the potential of a phylogenetic community approach to understanding food webs dynamics.

## Introduction

Predators can have strong top-down effects, both on community structure and ecosystem processes (Estes et al. 2011). The combined effect of predator species on communities is often stronger or weaker than that predicted from a study of those same species in isolation (Sih et al. 1998; Ives et al. 2005). These non-additive effects occur when predators interact with each other directly, or via their shared prey species. For example, predators feed directly on each other (intra-guild predation), consume the same prey (resource competition) or modify the behaviour of prey or other predator species (Sih et al. 1998; Nyström et al. 2001; Griswold and Lounibos 2006). These non-additive effects can be positive or negative. For example, prey may have an induced defense against one predator which increases (negative non-additive effect) or decreases (positive non-additive effect) the likelihood of consumption by a second predator. While there are many possible mechanisms underlying the effect of predator composition, we lack a means of predicting *a priori* the strength and direction of this effect on community structure and ecosystem function.

The phylogenetic relationships among predators could provide a framework for combining different approaches to studying predator-predator interactions, thus helping us make predictions about combined effects of predators. A phylogenetic approach to species interactions extends the measurement of species diversity to include the evolutionary relationships between species. Relatedness may be a proxy for ecological similarity; very similar species may compete strongly, and/or may interfere with each other while very different species may not be able to occur in the same patch. This approach was first used to interpret observations of community structure, as ecologists interpreted nonrandom phylogenetic structure (i.e.~under-or over-dispersion) as evidence for processes, such as habitat filtering or competition, which structure communities (Webb et al. 2002; Cavender-Bares et al. 2009). Recently, this approach has been applied to manipulative experiments. For example, the phylogenetic diversity of plant communities is a better predictor of productivity than either species richness or diversity (Cadotte et al. 2008; e.g. Cadotte et al. 2009; Godoy et al. 2014). In all cases, an implicit assumption is that increased phylogenetic distance is associated with increased ecological dissimilarity – either in the form of differences in species niches, interactions, or functional traits. When this is true, high phylogenetic diversity should lead to complementarity in resource use between species, resulting in increased ecosystem functioning (Srivastava et al. 2012).

Phylogenetic diversity may be a better predictor of species effects on ecosystem funcitioning than species identity alone. For example, studies of plants (Cadotte et al. 2008) have shown that ecosystem function is positively related to the phylogenetic diversity of plants. Although there have been many studies taking a phylogenetic approach to community ecology and although predators have large effects on many communities, the phylogenetic diversity of local predator assemblages has rarely been measured (Bersier and Kehrli 2008; Naisbit et al. 2011). Many studies of phylogeny and predator traits focus on whole clades, rather than local assemblages (e.g. *Anolis* lizards (Knouft et al. 2006), warblers (Böhning-Gaese et al. 2003), tree boas (Henderson et al. 2013) and wasps (Budriene and Budrys 2004)), making it difficult to connect these results to predator effects at the scale of a local community. These clade specific studies often find weak evidence for phylogenetic signal in ecologically relevant traits. In contrast, studies at the level of the whole biosphere (Bersier and Kehrli 2008; Gómez et al. 2010) demonstrate that related organisms often have similar interspecific interactions, i.e.~related predators often consume similar prey. At the local scale, only a few studies have examined how phylogeny may shape food webs (Rezende et al. 2009; Cagnolo et al. 2011); these observational studies found that models containing both relatedness (either from taxonomic rank or phylogenetic trees) and body size were better at predicting which predator-prey interactions occurred than models with body size alone. As observational studies, however, they cannot isolate if it is differences in predator distribution or diet that leads to a phylogenetic signal in predator-prey interactions, nor how these interactions affect the whole community.

Can phylogeny help us predict how predators will impact community composition and ecosystem functioning? Within a local community, the effect of predator species diversity will depend on three factors: how predators are distributed among habitats, how they interact with their prey, and how they interact with each other. To the extent that phylogenetic relationships are correlated with these three factors, they enable us to predict the impact of predator diversity on communities. For instance, phylogeny could constrain predator species co-occurrence if more distant relatives have more distinct fundamental niches, whereas close relatives are too similar to co-exist (Webb et al. 2002; Emerson and Gillespie 2008). When predators do co-occur, phylogeny may correlate with their feeding behavior, such that closely related predators consume similar prey. For example, diet overlap (shared prey species be-tween predators) will depend on the feeding traits and nutritional requirements of predators – both of which may be phylogenetically conserved. If this is the case, then predator as-semblages with higher phylogenetic diversity will show a greater range of prey consumed and therefore stronger top-down effects (Finke and Snyder 2008). In some cases, predator diets may extend to include other predators, leading to direct negative interactions such as intraguild predation, which may also have a phylogenetic signal (Pfennig 2000). To our knowledge, the relationship of phylogeny to predator distribution, diet, and intraguild inter-actions has never been investigated in a single study.

We tested for the effects of phylogenetic distance on the distribution, diet and interactions of predators living in a natural mesocosm: water reservoirs found inside bromeliad leaves. Bromeliads (Bromeliaceae) are flowering plants abundant in the Neotropics. Within this aquatic food web, damselfly larvae (e.g. *Leptagrion* spp., Odonata:Coenagrionidae) are important predators that dramatically reduce insect colonization (Hammill et al. 2015) and emergence (Starzomski et al. 2010), and increase nutrient cycling (Ngai and Srivas-tava 2006). In addition to damselfly larvae, other predators are also found in bromeliads, including large predaceous fly larvae (Diptera: Tabanidae) and predatory leeches (Hiru-dinae:Arhynchobdellida) (see Frank et al. (2009)). Many bromeliads contain water and trapped, terrestrial detritus which supplies nutrients for the bromeliad (Reich et al. 2003). The small size of these habitats permits direct manipulations of entire food webs, manipulations which would be difficult in most natural systems. Predators have been shown to have large top-down effects on ecosystem functions in bromeliads, including nitrogen uptake by the plant (Ngai and Srivastava 2006), detrital decomposition, and CO_2_ flux (Atwood et al. 2013; Atwood et al. 2014).

We tested for a relationship between the distribution, diet and ecosystem effect of predators and their phylogenetic distance using observations, lab feeding trials, and manipulative field experiments, respectively. We observed the distribution of predators between bromeliads by dissecting a sample of natural bromeliads. We quantified diet preferences in a series of no-choice feeding trials. We measured ecosystem-level effects with a manipulative experiment: we added predators to standardized bromeliad communities, adding either a single predator species or a pair of species of varying phylogenetic distance. In each approach, we test the hypothesis that the phylogenetic distance between predators determines the net impact of predator assemblages on the bromeliad commuinty:

1. *Distributional similarity*: We predict that closely related predators occur in the same habitat patch more frequently than less related predators. Alternatively, closely related species may never co-occur because of competitive exclusion.
2. *Diet similarity*: We predicted that closely related predators will eat similar prey at similar rates. Alternatively, closely related species may have evolved different diets to facilitate coexistence.
3. *Ecosystem-level effects*: We tested two sets of hypotheses about direct and indirect effects of predator combinations on ecosystems, predicting:

a. Closely related predators will have similar individual effects on the community. This will occur if related predators have similar trophic interactions (e.g. predation rate, diet similarity). Our single-species treatments allow us to assess the effect of each predator both on prey survival and on ecosystem functions.
b. Predator assemblages with higher phylogenetic diversity will have synergistic (greater than additive) effects on prey consumption and associated ecosystem functions. This will occur if phylogenetic distance correlates with increasing trait difference, and if this trait difference in turn results in niche complementarity. However, at the extreme, different predators may consume each other, thus creating antagonistic (less than additive) effects on prey consumption. By comparing treatments with pairs of predators to treatments that received each predator alone, we are able to estimate additive and non-additive effects.

## Methods

### Study Design

We used three empirical approaches to test the hypotheses outlined above. To test hypothesis 1 (distribution) we sampled bromeliads for predator species. To test hypothesis 2 (diet similarity), we conducted a series of laboratory feeding trials. Finally, we tested hypothesis 3 (similarity of community effect and interaction) with a field experiment in which predators were added to bromeliads containing standardized communities of prey. This experiment included both single species treatments and two species treatments; the latter were chosen to create the widest possible range of phylogenetic diversity.

We included phylogenetic information in our analyses of all three datasets. We obtained this phylogenetic information first from classification alone. Next we added information about the age of each node from “timetree.org”, an online database of published molecular time estimates (Hedges et al. 2006). The timetree online database collects information from multiple independent phylogenetic studies. These studies provide independent estimates of the age of the most recent common ancestor for two lineages. Lineages that diverged a long time ago have been dated by multiple studies; for such nodes we used the median age. All internal nodes were dated by at least one study, however data was unavailable for the youngest nodes (i.e. tips) of the tree. For these nodes, either a lack of taxonomic information (e.g. Tabanidae) or a lack of phylogenetic study (e.g. *Leptagrion*) prevented more information from being included. These branches were left unresolved (i.e., as polytomies) and were all assigned identical, arbitrary and short branch lengths (15 Mya). The result is a phylogeny that closely resembles the qualitative, taxonomy-based tree with which we began. Because the node ages between our major predators (leeches, tabanids and odonata) are so deep, variation among studies in the estimated age of these nodes was minor compared to the differences between them Our final tree is available at https://dx.doi.org/10.6084/m9.figshare.3980349.v1.

We conducted all three experiments in Parque Estadual da Ilha do Cardoso (25° 03’ S, 47° 53’ W), a 22.5 ha island of the south coast of São Paulo state, Brazil. We worked in a coastal forest (*restinga*) with an understory dominated by *Quesnelia arvensis* Mez. (Bromeliaceae). *Q. arvensis* is a large terrestrial bromeliad that catches and holds rainwater (phytotelmata), accumulating up to 2.8 L of rainwater in a single plant. Our observational survey found more than 47 species of macroinvertebrates in these aquatic communities (Romero and Srivastava 2010), in 25 bromeliads of various sizes. This diversity encompasses multiple trophic and functional groups. Filter feeders were entirely mosquito larvae (Diptera:Culicidae); detritivores include shredders (Diptera:Tipulidae, Trichoptera:Calamoceratidae), scrapers (Coleoptera:Scirtidae), and collectors (All Diptera:Chironomidae, Syrphidae, Psychodidae). All these species are prey for a diverse predator assemblage dominated by at least three species of damselfly larvae (*Leptagrion* spp., Odonata:Coenagrionidae), two species of horsefly larvae (Diptera:Tabanidae), and two species of leech (Arhynchobdellida). A lower percentage of predator biomass was composed of Dytiscid larvae (Coleoptera), midge larvae (Diptera: Ceratopogonidae) and chironomid larvae (Diptera: Tanypodinae).

## Data collection

### Distributional similarity

We asked whether closely related predators were found in the same bromeliads. In 2008, each bromeliad was dissected and washed to remove invertebrates. We passed this water through two sieves (150 and 850 μm), which removed particulate organic matter without losing any invertebrates. All invertebrates were counted and identified to the lowest taxonomic level possible. The body length of all individuals was measured when possible for small and medium-sized taxa (< 1cm final instar) and always for large-bodied taxa (> 1 cm final instar).

### Diet Similarity

To test whether related predators eat similar prey, we fed prey to predators in laboratory feeding trials. We conducted 314 feeding trials of 10 predator taxa and 14 prey taxa between March and April 2011. We included all potential predator-prey pairs present in the experiment (described below), and attempted to perform all other combinations whenever possible. However, due to the rarity of some taxa, many predator-prey pairs were not possible to assemble in the field; we tested 56 pairwise combinations. Most trials were replicated at least five times, but the number of replicates ranged from 1 to 11. To conduct the trials, we placed predators together with prey in a 50ml vial, with a stick for substrate. The only exception was the tabanid larvae, which we placed between two vertical surfaces to imitate the narrow space found in bromeliad leaf axils (their preferred microhabitat, necessary for successful feeding). Generally our trials contained a single predator and a single prey individual, except in the case of very small prey (*Elpidium* sp.) or predators (*Monopelopia* sp.), in which case we increased the density. We recorded whether prey was consumed after 24 hours. All feeding trial data is available at https://dx.doi.org/10.6084/m9.figshare.3978783.v1

### Community effect experiment

Our third hypothesis had two parts: (a) how do predator species differ in their individual effects on the invertebrate community composition (the number of surviving prey species) and ecosystem processes (rates of detrius consumption and nitrogen cycling) and (b) do predator combinations show non-additive effects on community and ecosystem processes, and do these non-additive effects increase or decrease with phylogenetic distance?

We tested effects of both single and multiple predator species on community responses with a manipulative experiment where identical prey communities were exposed to treatments of either a single predator, or pairs of predators representing increasing phylogenetic diversity. In this experiment we focused on the four most abundant large predators found in the com-munity: *Leptagrion andromache* and *Leptagrion elongatum* (Odonata: Coenagrionidae), a predatory Tabanid fly (Diptera:Tabanidae:*Stibasoma* sp.) and a predatory leech. We com-bined these species in eight treatments: predator-free control (no predators), each of the four predator species alone (3a) and pairs of predator species chosen to maximize variation in phylogenetic distance (3b). Specifically, these pairs were: two congeneric damselflies (*Lep-tagrion andromache* and *Leptagrion elongatum*), two insects (*L. elongatum* and *Stibasoma*), and two invertebrates (*L. elongatum* and a predatory leech). We used five replicate bromeli-ads for each of these 8 treatments (8 treatments, n=5). This experiment, therefore, allows the estimation of the effect of each predator species (single-species treatments), as well as the detection of non-additive effects in predator combinations.

We created bromeliad communities that were as similar as possible to each other, and also to the average composition of a bromeliad. In February 2011 we collected bromeliads with a volume between 90 and 200ml, thoroughly washed the plants to remove organisms and detritus, and soaked them for 12 hours in a tub of water. We then hung all bromeliads for 48 hours to dry. This procedure was intended to remove all existing macroinvertebrates; one bromeliad dissected afterwards contained no insects (a similar technique was used by Romero and Srivastava (2010)). We simulated natural detritus inputs from the canopy by adding a standard mass of dried leaves of the species *Plinia cauliflora* (Jabuticaba, Myrtaceae; a common Brazilian tree; 1.5g bromeliad ^−1^ ± 0.02, mean ± sd). In order to track the effects of detrital decomposition on bromeliad N cycling, we enriched these leaves with ^15^N by fertilizing five plants with 40ml pot^−1^ day^−1^ of 5g L^−1^ ammonium sulphate containing 10% atom excess of ^15^N. After 21 days we then collected *P. cauliflora* leaves, air-dried until constant weight, and then soaked them for three days. This procedure removes excess nutrients from the artificial fertilization. Because some of our prey species consume fine detritus, not coarse, we also added a standard amount of dried fine detritus to our bromeliads (0.23g bromeliad ^−1^ ± 0.02). This fine detritus originated from detrital material betwee 150 and 850 micrometers in size obtained from unmanipulated bromeliads and oven-dried.

Each bromeliad was stocked with a representative insect community (See supplementary material). The densities of each prey taxon were calculated from the observational dataset (Hypothesis 1), using data from bromeliads of similar size to those in our experiment. We ran this experiment in two temporal blocks for logistical reasons: three complete replicates of all treatments were set up on 20 February 2011, and two on 08 March 2011. We first placed the prey species into the bromeliad, allowed two days for the prey to adjust, then added predators. After 26 days from the beginning of each block, we added the same prey community a second time to simulate the continuous oviposition that characterizes the system. We concluded the experiment 43 days from the first addition of prey (20 April 2011). Throughout the experiment, all bromeliads were enclosed with a mesh cage topped with a malaise trap and checked daily for emergence of adults. At the end of the experiment we completely dissected our bromeliads, collecting all invertebrates and detritus remaining inside.

We used a substitutive design, maintaining the same predator metabolic capacity in all replicates (see below). In a substitutive experiment, all experimental units receive the same “amount” of predators – usually standardized by abundance – and only species composition varies. However, when species differ substantially in body size - as in this experiment - abun-dance does not standardize the their effects on the community. We chose to standardize using metabolic capacity instead (after Srivastava (2009)). Integrating the allometric relationship between body size and feeding rate (Brown et al. 2004; Wilby et al. 2005) over all individuals of a species allows estimates of “metabolic capacity”, or the potential energy requirements of a species (Srivastava and Bell 2009). Metabolic capacity is equal to individual body mass raised to the power of 0.69 (an invertebrate-specific exponent determined by Peters (1986) for invertebrates and confirmed by Chown et al, (2007)); this reflects the nonlinear relationship between feeding rate and body size across many invertebrate taxa.

To quantify the effect of predators on ecosystem function, at the end of the experiment we measured five community and ecosystem response variables: decomposition of coarse detritus, production of fine particulate organic matter (FPOM), bromeliad growth, uptake of detrital nitrogen into bromeliad tissue, and survival of invertebrate prey (emerged adults + surviving larvae). We measured decomposition by passing the bromeliad water through a 850 μm sieve, collecting the retained detritus and determining the mass of this detritus after oven-drying it at approximately 70°C. We measured the production of FPOM by taking the remaining liquid and filtering it on pre-weighed coffee filters, which were then dried and reweighed. We measured bromeliad growth as the average increase in length of five leaves per plant. We tracked the uptake of labeled detrital nitrogen by analyzing the isotopic composition of the three innermost (closest to meristem) bromeliad leaves at the end of the experiment. These analyses were performed at the Stable Isotope Facility laboratory (UC Davis, CA, USA) using continuous flow isotope ratio mass spectrometer (20-20 mass spectrometer; PDZ Europa, Sandbach, England) after sample combustion to N_2_ at 1000°C by an on-line elemental analyzer (PDZ Europa ANCA GSL). Finally, we quantified the species composition and survivorship of invertebrate prey by combining counts of emerging adult insects and surviving larvae. All experimental data is available at https://dx.doi.org/10.6084/m9.figshare.3983964.

## Data analysis

We quantified the effect of phylogenetic distance on each of distributional (Hypothesis 1) and diet (Hypothesis 2) similarity. First, we calculated phylogenetic distance between each pair of species. We then evaluated both distributional and diet similarity between predators using Pianka’s index of niche overlap (Pianka 1974):

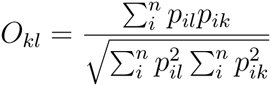

For each pair of predators, *p*_*ik*_ and *p*_*il*_ represent the preference of predator *k* or *l* for resource or habitat *i*. The value *O*_*kl*_ represents similarity (in our case, in either distribution or diet) and ranges from 0 (complete dissimilarity) to 1 (complete similarity). The *n* resources represent the different habitats surveyed for Hypothesis 1 (distributional similarity), or the different prey species assayed for Hypothesis 2 (diet similarity). Preference (*p*_*ik*_) represents the proportion of a predator’s total metabolic capacity found in a particular bromeliad (Hypothesis 1); or the proportion of feeding trials in which it ate a particular prey (Hypothesis 2). We then compared these effects via a Mantel test, to check for overall correlation between the phylogenetic distance matrix and dissimilarity in either predator distribution or diet preferences.

We divided the analysis of the manipulative experiment into three parts: quantifying the effect of phylogenetic distance on prey community similarity, on community and ecosystem responses, and on non-additive effects of predator combinations. First, we compared the four treatments with single predator species by calculating the similarity in species composition (Pianka’s index) between surviving prey communities and relating this to the phylogenetic distance between predators with a linear regression. If predator feeding choices are phylogenetically conserved, then diet similarity will decline with increasing phylogenetic distance.

Second, we measured five community and ecosystem responses, testing in turn the effect of predator presence, number, species identity, and finally phylogenetic diversity. To test for an effect of predator presence, we compared the control treatment (predators absent) with the mean responses of all seven treatments that did contain predators. To test for an effect of predator species number (one or two predators), we compared the means of all single-species treatments with the means of all two-species treatments. To test for an effect of predator identity, we compared all four single-species treatments. Finally, to test for an effect of predator combinations we compared all two-species treatments (3 pairs total). We analyzed each of these of these orthogonal comparisons with one-way ANOVA.

In our third and final analysis, we quantified the non-additive effect of predator species on our responses. We calculated this effect as the difference between the response in bromeliads with both predator species (n=5) and the mean response in bromeliads with either one of these two predator species (n=5 for each predator species). We generated bootstrap confidence intervals for these non-additive effects; confidence intervals that do not overlap zero indicate a significant non-additive effect of a predator combination. We used R version 3.2.0 (R Core Team 2015) for all calculations, and two packages: picante (Kembel et al. 2010) for calculating phylogenetic distances matrices, and vegan (Oksanen et al. 2015) for distance metrics. All the code documenting our analyses is archived at http://dx.doi.org/10.5281/zenodo.16805

## Results

### Hypothesis 1: similarity in distribution

We did not find any significant relationship between the co-occurence of a pair of predators in bromeliads (measured as Pianka’s index of niche overlap) and the phylogenetic distance between the two predators. A Mantel test found no evidence of correlation between differences among predators in habitat use, and phylogenetic distance (correlation −0.18, p = 0.82, 999 permutations). This indicates that all 14 predator species have roughly similar habitat distri-butions – indeed, we often found multiple predator species co-occurring in the same bromeli-ads (mean 4.45 ± 2.8 predator species per plant). We were able to sample a wide range of phylogenetic relatedness, including two groups of congenerics – two species of *Bezzia* sp. (Diptera:Ceratopogonidae) and three species of *Leptagrion* sp. (Odonata:Coenagrionidae). There were also two groups of confamilials – three species of Tabanidae and two species of Empididae, all Diptera. Deeper divisions were also present: three families of Diptera were represented by a single predator species each (Dolichopodidae, Corethrellidae and Chironomidae) and the deepest taxonomic divide was between all insects present and the predatory leeches (Arhynchobdellida:Hirudinidae).

### Hypothesis 2: Similarity in diet

Overall, predators were remarkably similar in their diets, reflecting the broad generalist diets of most predators (Fig. 1b). Although diet similarity appears to decline slightly with phylogenetic distance between predators, this effect disappears once we correct for non-independence of predator pairs with a Mantel test (correlation −0.27, p = 0.88, 999 permutations).

**Figure 1:**
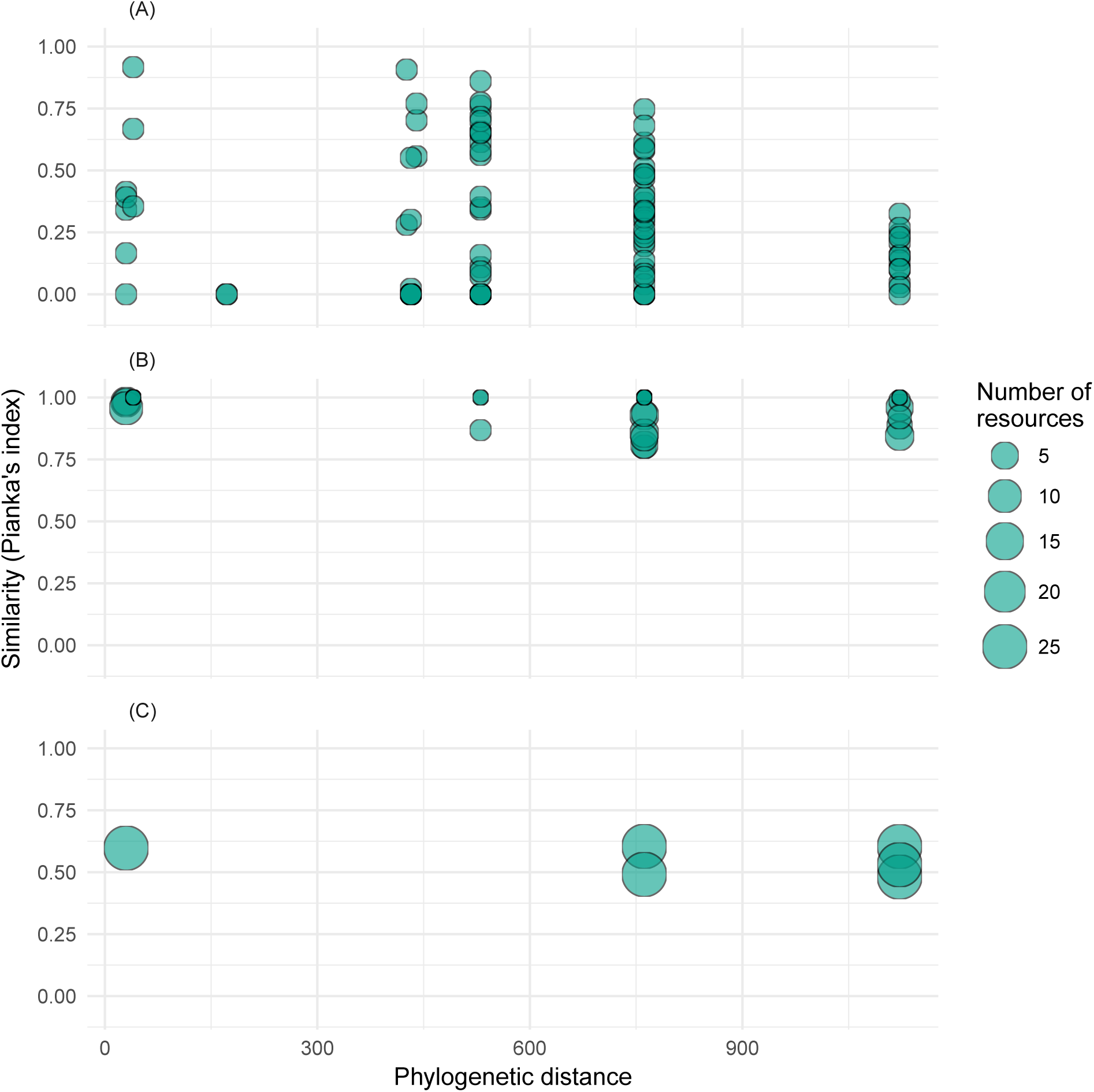
Phylogenetic distance between predators as a predictor of niche overlap among predators and impacts on prey composition. Our measures of niche overlap were: (a) distribution among bromeliads and (b) diet preferences. We also show the effect of phylogenetic distance between predators on (c) community dissimilarity of surviving prey (Bray-Curtis dissimilarity). We measured distributional similarity (a) by counting all predators in 25 bromeliads, estimating their total metabolic capacity, and calculating niche overlap (Pi-anka’s index) among all pairs of species. We measured diet preferences (b) for a subset of these predators by offering them various prey in no-choice trials, and again calculated niche overlap among them. Finally, we measured community composition of surviving prey (c) at the end of an experiment in which predators were placed in bromeliads with standardized prey communities. For (a) and (b) we used Pianka’s index of niche overlap (1 = complete niche overlap) and tested various nonlinear and linear models (see Appendix) of the relationship between this index and phylogenetic distance. Solid lines show significant model ft, and dashed lines show bootstrap 95% quantiles.

### Hypothesis 3: similarity in top-down effects

We analyzed our five univariate response variables from the manipulative experiment by di-viding them into four separate and orthogonal tests: predator presence, predator number, predator species identity, and increasing predator phylogenetic diversity. Across all four tests, we saw the strongest responses in terms of total prey survivorship (Table 1). Prey sur-vivorship was halved when predators were present (Figure 2a, Table 1). Although predator species differed in their individual effects on the composition of the surviving prey com-munity, this difference was unrelated to the phylogenetic distance between predator species (Fig 1c, F_1,4_=0.71, p=0.45, distance measured as Bray-Curtis dissimilarity). Although single predator species had similar effects on prey survivorship (Figure 2c, Table 1), combinations of predators with higher phylogenetic diversity showed a significant increase in total prey survivorship (Fig 2d). That is, more phylogenetically diverse pairs of predators caused less prey mortality. Interestingly, these antagonistic effects on prey surviorship did not result in a change in the processing of detritus (measured either as reduction in coarse detritus or production of fine detritus), bromeliad growth or nitrogen cycling (Table 1).

**Table 1:** Predator diversity effects on community and ecosystem variables. We measured five community-level variables: total prey survival (both emerged adults and surviving larvae; see Fig. 2 and 3), the breakdown of coarse detritus (decomposition), the production of fine particulate organic matter (FPOM), the cycling of nitrogen from detritus to bromeliad tissue, and the growth of the bromeliad itself. We contrast treatments in our experimental design in four orthogonal ways: comparing treatments with predators to those without (“Predator Presence”), contrasting predator species (“Identity”), comparing predator communities of 1 or 2 species (“Richness”), and considering the effects of phylogenetic distance between predators (“Pairwise PD”). Values are slope ± standard error and = p < 0.05

| Response | Predator Presence | Identity | Richness | Pairwise PD |
| --- | --- | --- | --- | --- |
| Total prey survival | $-7.37 \pm 2.45$; $F_{1,10} = 9.07^*$ | $2.00 \pm 2.07$; $F_{3,16} = 0.60$ | $2.05 \pm 1.46$; $F_{1,5} = 1.96$ | $0.01 \pm 0.00$; $F_{1,13} = 7.64^*$ |
| Decomposition (g) | $0.01 \pm 0.02$; $F_{1,10} = 0.47$ | $-0.01 \pm 0.03$; $F_{3,15} = 1.29$ | $-0.01 \pm 0.02$; $F_{1,5} = 0.21$ | $0.00 \pm 0.00$; $F_{1,13} = 0.40$ |
| FPOM (g) | $-0.06 \pm 0.09$; $F_{1,10} = 0.46$ | $-0.06 \pm 0.11$; $F_{3,15} = 0.28$ | $0.18 \pm 0.07$; $F_{1,5} = 6.19$ | $-0.00 \pm 0.00$; $F_{1,13} = 1.45$ |
| Bromeliad growth | $-0.79 \pm 1.10$; $F_{1,10} = 0.51$ | $-1.08 \pm 1.62$; $F_{3,16} = 0.96$ | $0.59 \pm 0.84$; $F_{1,5} = 0.49$ | $0.00 \pm 0.00$; $F_{1,12} = 1.29$ |
| nitrogen cycling | $-5.69 \pm 4.03$; $F_{1,10} = 2.00$ | $-0.22 \pm 8.66$; $F_{3,16} = 1.84$ | $3.97 \pm 5.63$; $F_{1,5} = 0.50$ | $-0.00 \pm 0.01$; $F_{1,13} = 0.15$ |

**Figure 2:**
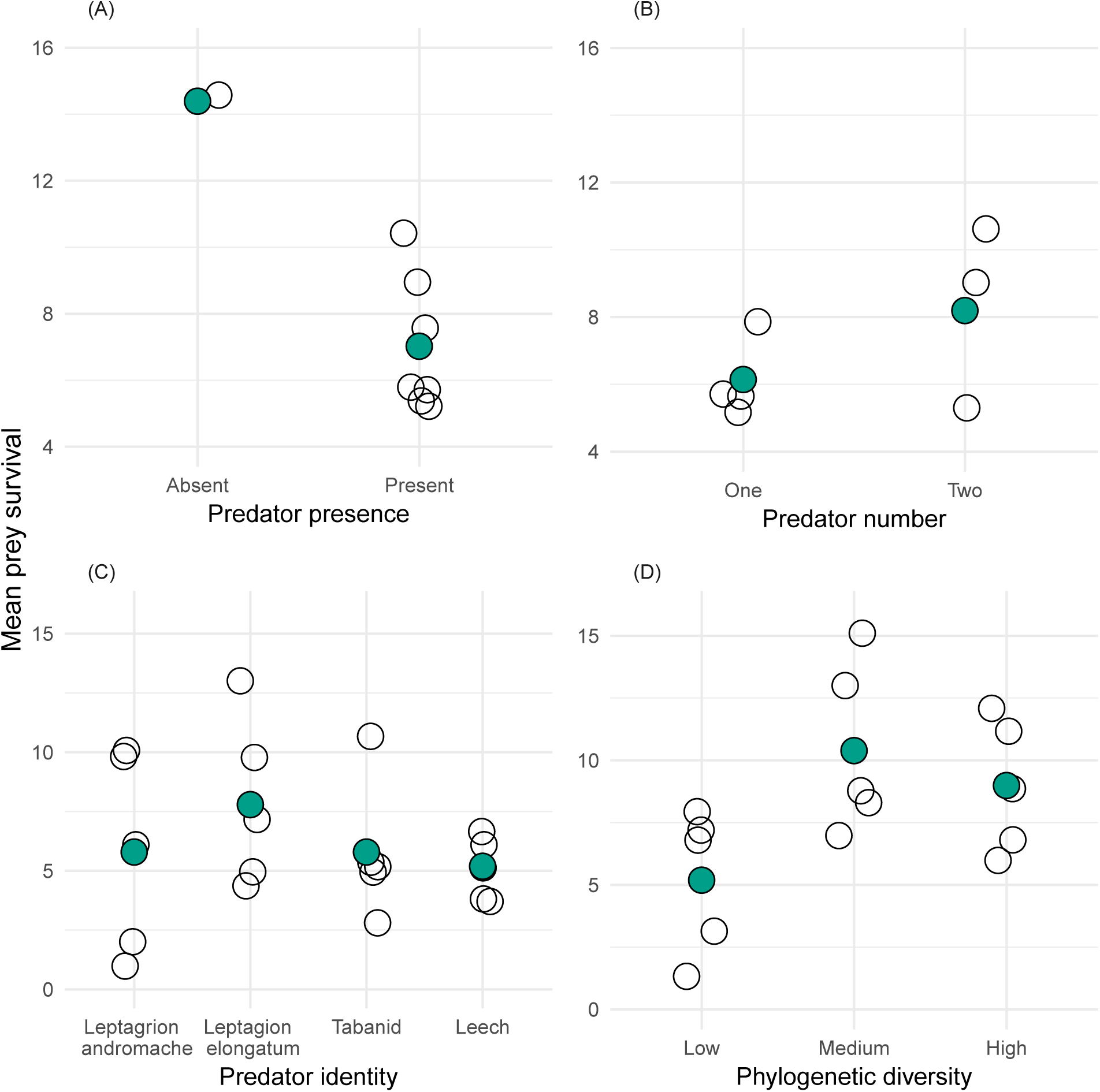
Orthogonal comparisons of the effect of predators on prey survival. We show the effects of predator presence (a), and then within predator present treatments the effects of predator species number (b). Within treatments with one predator species, we show effects of predator identity (c). Within treatments with two predator species, we show the effect of increasing phylogenetic diversity (d, arranged in order of increasing phylogenetic distance: Low = *L. andromache* + *L. elongatum*, Medium = *L. elongatum* + tabanid, High = *L. elongatum* + leech). Shaded dots represent grand means for each group; unshaded dots are either treatment means (2a and 2b, n = 5) or individual bromeliads (2c and 2d). Points are jittered horizontally slightly to reveal all datapoints.

We tested for non-additive effects of predator phylogenetic diversity with bootstrap confi-dence intervals. When we compared the actual effects of predator combinations with those expected from the mean of each single-species treatment, we found that predator pairs with the greatest phylogenetic diversity had the highest prey survival (Table 1). Whereas effects of *L. andromache* and *L. elongatum* in combination were quite similar to the effect of either alone, when *L. elongatum* was placed in the same plant as either a *Stibasoma* larva or leeches, on average five more prey individuals (18% of total prey community) survived till the end of the experiment (Fig 3; Tabanid, p = 0.016, Leech, p = 0.016). Once again, this effect on invertebrate density did not in turn create a significant difference in the ecosystem function variables.

**Figure 3:**
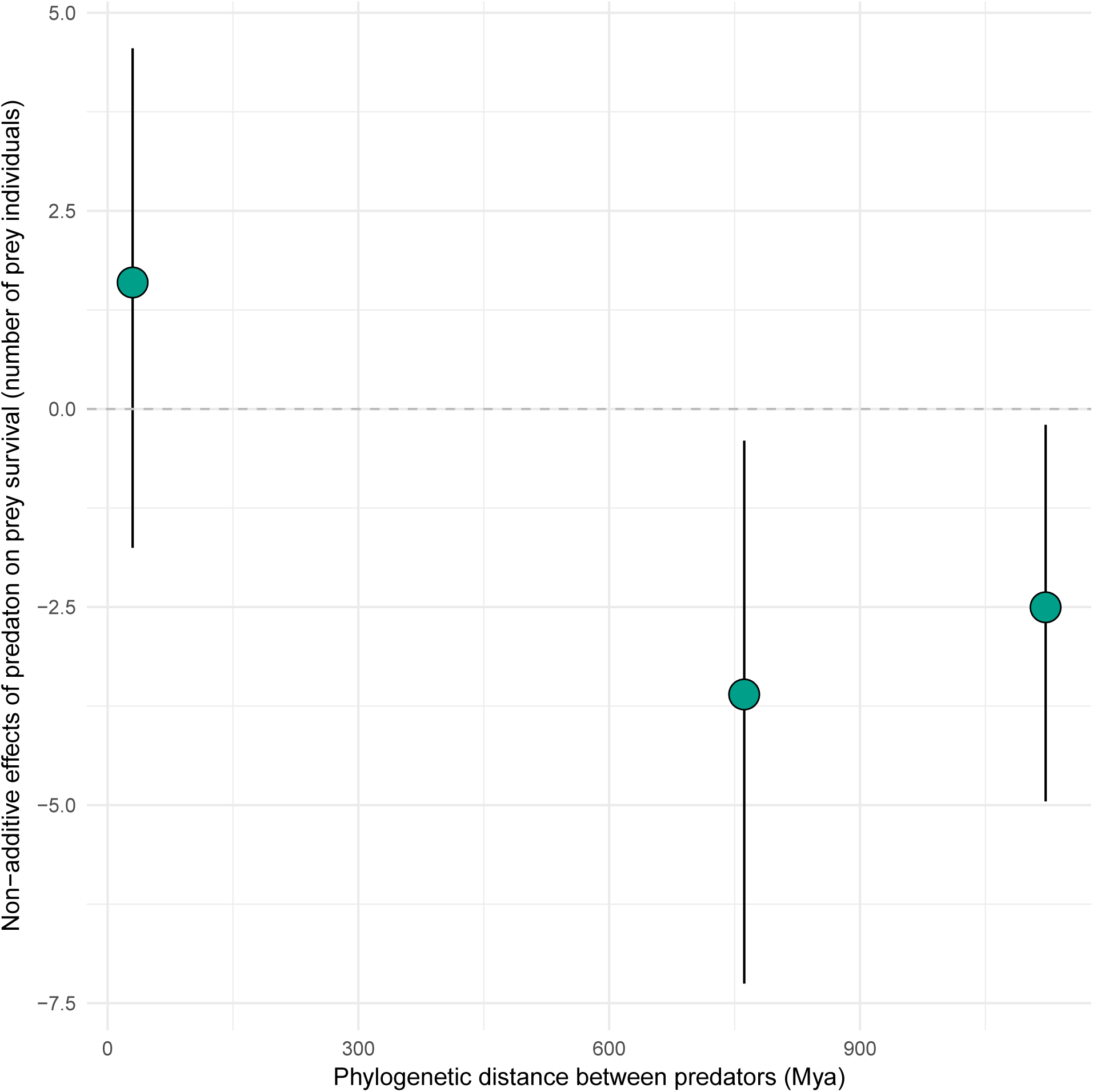
Non-additive effects of predator combinations on prey decrease with increasing phylogenetic distance between predators. A difference of 0 indicates that two-predator treat-ments resulted in no more prey mortality than would be expected from simply averaging single-predator treatments. A negative difference indicates that two-predator treatments resulted in less mortality than expected. Error bars represent bootstrap 95% confidence intervals.

## Discussion

We found that the phylogenetic distance between predators had variable importance in the bromeliad system. The phylogenetic distance between predators was unrelated to their co-occurrence (Hypothesis 1). The phylogenetic distance between predator species was also unrelated to diet overlap, although there was a tendency of diet overlap to decrease by about 20% over the observed range of phylogenetic distance (Hypothesis 2). Perhaps as a consequence of diet similarity, the phylogenetic distance between predators could not predict their individual effects on prey composition or survival (Hypothesis 3a). However, greater phylogenetic diversity caused an increase in prey survival (i.e. a decrease in predation); phylogenetically distant pairs of predators that co-occurred in bromeliads had less impact on prey than expected from their performance in isolation (Hypothesis 3b). We examine each of these main results in turn.

### Phylogenetic distance and similarity in distribution

Phylogenetic distance between predators did not explain overlap in habitat distribution. This similarity in distribution could be caused by two processes: low habitat variability among bromeliads, or low variability in preference of predators for different habitats. Bromeliads at this site vary widely in abiotic conditions, size, detritus amount and prey community; therefore it seems unlikely that low patch variation explains the lack of pattern. It appears instead that predators do not possess any strong phylogenetically-conserved preferences for different habitat characteristics, showing instead very generalist habitat preferences. This is not surprising, given that these organisms live in small, fluctuation-prone habitats. As a group, predatory invertebrates in bromeliads do not show more sensitivity to bromeliad size or drought than other invertebrates (Amundrud and Srivastava 2015). The co-occurrence of predator species within bromeliads suggests that antagonistic interactions among predators do not limit species distributions. Additionally, it appears that predator species are able to co-occur in many different combinations, creating a range of phylogenetic diversities within bromeliads. This suggests that the range of phylogenetic diversity we tested in our experiment was realistic.

### Phylogenetic distance and similarity in diet

There was no significant relationship between phylogenetic distance and overlap in diet as measured by laboratory feeding trials. n part, this reflects the ability of many predator species to consume a range of prey. However, predator species still showed some differences in prey preference. For example, damselflies are visual predators that engulf prey whole using specialized mouthparts; they are gape-limited and cannot eat prey that are too large. Leeches, in contrast, lack eyes but are able to pierce prey and consume them without swallowing. Damselflies showed a much stronger preference for culicid larvae than did leeches, whereas leeches were slightly better able to kill and consume scirtids. Culicid larvae are free swimming in the water column, and are therefore easily captured by engulfing predators, whereas scirtid larvae crawl on surfaces and are difficult to remove. Despite these modest differences between predator species in diet, such differences appeared largely unstructured by phylogeny. Other studies have also suggested that predator functional traits are more important than phylogeny *per se* to a predator’s diet: Moody (1993) found that unrelated decapod species which were morphologically similar were also functionally similar. Similarly, Rezende et al. (2009) found that both body size and phylogeny determined the food web “compartment” (shared predator-prey interactions) of predators in a marine foodweb.

### Phylogenetic distance and non-additive effects

We found that the presence of predators reduced prey survival, but that this reduction was less for phylogenetically-diverse combinations of predators. This was contrary to our hypothesis that more distant predators would show an increase in prey capture via niche complementarity. *L. andromache* did not produce an antagonistic (i.e. less than additive) effect in combination with *L. elongatum*, whereas the two more phylogenetically diverse combinations (*L. elongatum* with the Tabanid or leech) did. *Leptagrion* species may not distinguish between conspecifics and congenerics. In predicting a synergistic non-additive effect of predators, we were imagining an outcome much like those reported by Nilsson et al. (2006); they found that stoneflies caused prey to move into habitats where fish predators could consume them, increasing total predation (a synergistic effect, caused by a phylogenetically distinct predator). Our results are more consistent with those of Finke and Denno (2005), who found that combinations with two insect predators had a higher per-capita effect on leafhopper prey than combinations with an insect and a spider. That is, more phylogenetically diverse combinations of predators showed less predation on lower trophic levels.

When *L. elongatum* occurred with more distantly related predators, prey survivorship was greater than expected. This non-additive effect may have been due to a reduction in preda-tion by odonates in the presence of non-odonate predators. Odonates have been shown to be sensitive to chemical cues (Barry and Roberts 2014) or tactile cues (Atwood et al. 2014) of potential predators, which causes a decrease in feeding rate. For example, a different species of bromeliad damselfly – *Mecistogaster modesta* Selys – reduces predation when it is housed with Dytiscid adults (Atwood et al. 2014). If there is a phylogenetic signal to the chemical cues released by predators, individuals of one species might be unable to distinguish close relatives (congenerics in our case) from conspecifics. One limitation of our approach is that all phylogenetic diversity treatments contained one species in common, *Leptagrion elonga-tum*. It is possible that this species is more sensitive to the presence of other predators, and therefore shows a larger effect in combination than would other species in this community. However, this is the most common predator in this community and our results indicate that its top-down effects are likely to be frequently reduced by the presence of other predators.

In our experiment, we did not see any effect of predator presence, nor of increasing preda-tor phylogenetic diversity, on ecosystem function (defined here as nitrogen cycling, detritus decomposition and bromeliad growth). This was contrary to our predictions based on previ-ous studies from rainforest bromeliads, which found that adding predators to a community increased nitrogen cycling and reduced detrital decomposition (Ngai and Srivastava 2006; Srivastava and Bell 2009). While we did observe substantial consumption of detritivorous prey by predators, the resulting reductions in detritivore density did not cause differences in either the decomposition of detritus or the uptake of detrital nitrogen into bromeliad leaf tissue. These differences between our results and those from rainforests may be due to leaf traits of the *restinga* vegetation. In *restinga* vegetation, leaves are generally extremely tough and waxy, whereas in rainforests, leaves tend to be softer – with the result that, in *restinga*, invertebrates are unable to consume leaves directly. Several lines of evidence support this assertion. Romero and Srivastava (2010) studied the effects of the spider *Corinna demersa* (Corinnidae) on bromeliad ecosystems. This spider has no effect on the composition of detritivore communities, nor on decomposition rates, but increases nitrogen content in bromeliads, probably by depositing feces or the carcasses of terrestrial prey. This indicates that *restinga* bromeliads may derive less of their nitrogen from terrestrial detritus, but may benefit more from terrestrial inputs. A separate experiment (GQ Romero, pers comm) supports the hypothesis that lower decomposition in *restinga* is due to plant traits. This second experiment contrasted decomposition caused by invertebrates and bacteria with that caused by bacteria alone (by comparing bagged detritus enclosed in coarse vs fine mesh). The experiment used two species of detritus: leaves from a rainforest tree, and leaves from a *restinga* tree. Invertebrates only caused an increase in decomposition for the rainforest tree, not the *restinga* tree.

In most natural communities, multiple predator species co-occur and often simultaneously affect prey species. This study is one of the first to examine how phylogenetic diversity of a guild of predators affects both food web structure and ecosystem functioning. By combining an observational study, laboratory trials, and a field experiment that controlled number and phylogenetic diversity of predators we have shown that phylogenetic relatedness of species can help predict food web responses.

Previous studies have usually addressed this question in the context of species that only compete for resources, typically plants that compete for nutrients and water (Cavender-Bares et al. 2009). The predators in our system not only compete for prey, but also have the potential for intraguild predation. This adds a new way in which phylogenetic diversity can affect food webs and ecosystems. Phylogenetically distant predators may be more likely to prey on each other, either because injury is less likely when species differ in size and morphological defenses or, as suggested by Pfennig (2000), because the risk of disease transmission is less. If the risk of intraguild predation increases with predator phylogenetic diversity, this may counteract any ecosystem effects of diminished competition. When this is the case, increasing phylogenetic diversity may reduce overall predation rates, because predators fear intraguild predation from distantly-related predators, and simultaneously increase predation rates, because predators overlap less in prey preferences or in hunting mode. The net effects of these processes will be difficult to predict without detailed experiments like those that we report here.

Our results suggest that phylogenetic relationships among organisms at higher trophic levels may have more complex ecosystem consequences than when only a single, lower trophic level is considered. In order to apply phylogenetic community ecology to food webs, we will need to consider a broader suite of potential interactions between species and extend our theoretical framework beyond simple niche complementarity (Srivastava et al. 2012). However, this is a worthwhile goal. An approach based on phylogenetic diversity offers an organizing framework around which to compare diverse datasets on the distribution, trophic interactions and combined effect of multiple predator species, and to predict the top-down effect of diverse predator assemblages.

## Acknowledgements

We thank Aline Nishi, Robin LeCraw, Alathea Letaw, Gustavo Cauê Piccoli and Tiago Bernabé for support during field work. We also gratefully acknowledge the support of the Parque Estadual da Ilha do Cardoso and its residents, especially Eduardo Pereira. AAMM was supported by and NSERC CGS D scholarship and a Micheal Smith Foreign Study Supplement during this work. GQR is grateful to the Brazilian Council for Research, Development and Innovation (CNPq) and São Paulo Research Foundation (FAPESP) for research fund and productivity grant. DSS was supported by a NSERC Discovery grants and an E.W.R. Steacie Memorial fellowship.

